# Genome-wide identification of novel sRNAs in *Streptococcus mutans*

**DOI:** 10.1101/2021.11.16.468913

**Authors:** Madeline C Krieger, Justin Merritt, Rahul Raghavan

## Abstract

*Streptococcus mutans* is a major pathobiont involved in the development of dental caries. Its ability to utilize numerous sugars and to effectively respond to environmental stress promotes *S. mutans* proliferation in oral biofilms. Because of their quick action and low energetic cost, non-coding small RNAs (sRNAs) represent an ideal mode of gene regulation in stress response networks, yet their roles in oral pathogens have remained largely unexplored. We identified 15 novel sRNAs in *S. mutans* and show that they respond to four stress-inducing conditions commonly encountered by the pathogen in human mouth: sugar-phosphate stress, hydrogen peroxide exposure, high temperature, and low pH. To better understand the role of sRNAs in *S. mutans*, we further explored the function of the novel sRNA, SmsR4. Our data demonstrate that SmsR4 regulates the EIIA component of the sorbitol phosphotransferase system, which transports and phosphorylates the sugar alcohol sorbitol. The fine-tuning of EIIA availability by SmsR4 likely promotes *S. mutans* growth while using sorbitol as the main carbon source. Our work lays a foundation for understanding the role of sRNAs in regulating gene expression in stress response networks in *S. mutans* and highlights the importance of the underexplored phenomenon of posttranscriptional gene regulation in oral bacteria.

**IMPORTANCE:** Small RNAs (sRNAs) are important gene regulators in bacteria, but the identities and functions of sRNAs in *Streptococcus mutans*, the principal bacterium involved in the formation of dental caries, are unknown. In this study, we identified 15 putative sRNAs in *S. mutans* and show that they respond to four common stress-inducing conditions present in human mouth: sugar-phosphate stress, hydrogen peroxide exposure, high temperature, and low pH. We further show that the novel sRNA SmsR4 likely modulates sorbitol transport into the cell by regulating SMU_313 mRNA, which encodes the EIIA subunit of the sorbitol phosphotransferase system. Gaining a better understanding of sRNA-based gene regulation may provide new opportunities to develop specific inhibitors of *S. mutans* growth, thereby improving oral health.

## INTRODUCTION

The gram-positive oral pathogen *Streptococcus mutans* plays a principal role in the formation of dental caries and is often considered to be the primary causative agent of the disease (1–3). Central to *S. mutans* cariogenicity is its ability to ferment a wide variety of sugars, resulting in the formation of acidic microenvironments that drive the decline of commensal species and the proliferation of aciduric bacteria (4). One factor that allows *S. mutans* to metabolize a variety of carbon sources is the presence of 14 phosphotransferase systems (PTSs) (5, 6). PTSs act by transporting carbohydrates across the cell membrane and immediately phosphorylating them for intracellular retention and entry into glycolysis (7). While glucose is the preferred carbon source for *S. mutans,* other PTSs, including the sorbitol (glucitol) PTS, are inducible in its absence.

Sorbitol is a sugar-alcohol found naturally in many fruits and is a popular low-calorie anti-cariogenic sweetener (7, 8). The anticariogenic action of sorbitol is due to the relatively low amounts of acid produced during sorbitol fermentation by *S. mutans* compared to that of glucose or sucrose metabolism (8, 9).

The success of *S. mutans* in the oral cavity is enhanced not only by proficient sugar utilization but also by its ability to quickly respond to rapidly changing environmental conditions and multiple stressors (10). Small RNAs (sRNAs) are non-coding transcripts that typically bind to their target mRNAs to regulate gene expression (11). Due to their low energetic cost, fast action, and capacity for co-degradation with mRNA targets, sRNAs are an ideal mode of posttranscriptional control when a bacterium is experiencing environmental stress (12). Surprisingly little is known about sRNA-based regulation in *S. mutans* and there have been no functional analyses of sRNA utility in *S. mutans* to date. In this study, we identified 15 putative sRNAs in *S. mutans* that respond to multiple stress conditions and describe a novel sRNA that promotes bacterial growth in the presence of sorbitol.

## RESULTS

### Identification and validation of novel sRNAs

We used an RNA-seq-based approach that we developed previously (13–15) to identify 15 novel sRNAs — named SmsR for “*S. mutans* small RNA” — expressed in *S. mutans* during exponential (OD_600_ 0.5-0.6) growth in Brain Heart Infusion (BHI) broth (**Figure 1**). Nine of the novel sRNAs overlapped with candidates predicted by a previous genome-wide scan and a search against the Rfam database identified three of the sRNAs as RNase P, 6S RNA, and tmRNA (16, 17) (**Table 1**). All candidate sRNAs have putative -10 promoter sites, and 10 have predicted rho-independent terminators at their 3′ ends (**Table 2**), indicating these as authentic sRNAs. We further verified all 15 putative sRNA transcripts using Northern blot (**Figure 1**). Interestingly, in a few cases, higher molecular weight bands in addition to the sRNAs were observed, presumably because the sRNAs were cleaved from longer transcripts, as observed in other bacteria (18–23).

**Figures 1A and 1B.**
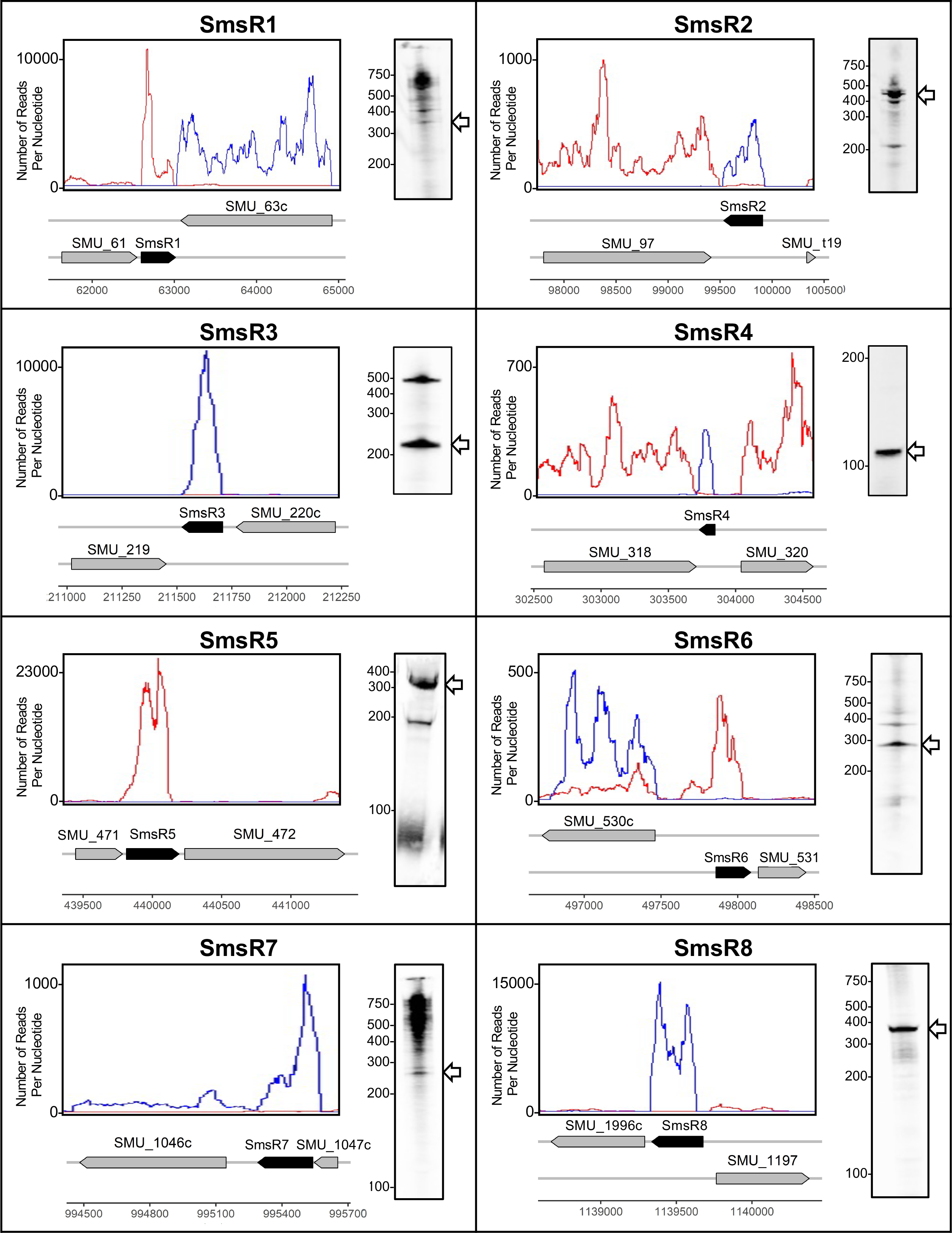
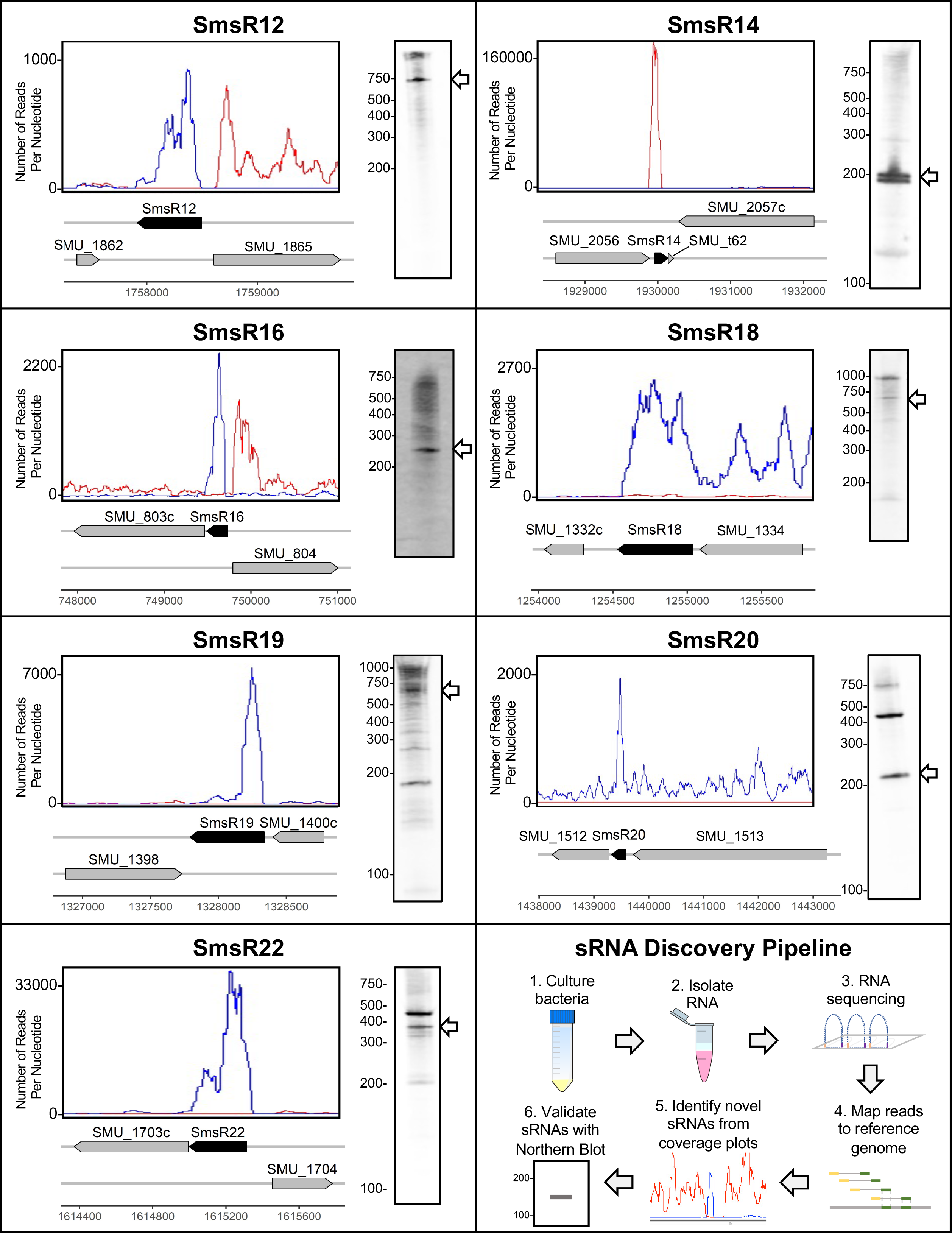
Expression profiles of novel sRNAs in *S. mutans*. RNA-seq reads mapped to forward (red) and reverse (blue) strands of the *S. mutans* UA159 genome are shown. Y-axes denote the number of RNA-seq reads mapped to each nucleotide. Genomic locations of novel sRNA genes (black arrows) and flanking genes (grey arrows) along with their nucleotide positions are shown below the coverage plots. Northern blot performed for each sRNA is shown to the right. White arrows denote estimated sizes of sRNAs (**Table 1**). (Note: intensity between each blot is not comparable as exposure times differed between experiments.)

**Table 1.** Novel sRNAs in S. mutans.

| sRNA<sup>a</sup> | Rfam annotation<sup>b</sup> | Previous prediction<sup>c</sup> | Bordering gene (left) | sRNA left bound<sup>d</sup> | sRNA right bound<sup>d</sup> | Bordering gene (right) | Strand | Predicted size (nt) |
| --- | --- | --- | --- | --- | --- | --- | --- | --- |
| SmsR1 |  |  | SMU_61 | 62597 | 62953 | SMU_63c | F | 357 |
| SmsR2 |  | psRNA-31 | SMU_97 | 99488 | 99909 | SMU_t19 | R | 422 |
| SmsR3 |  | psRNA-54 | SMU_219 | 211498 | 211709 | SMU_220c | R | 212 |
| SmsR4 |  | psRNA-62 | SMU_318 | 303748 | 303857 | SMU_320 | R | 110 |
| SmsR5 | RNase P | psRNA-73 | SMU_471 | 439833 | 440221 | SMU_472 | F | 389 |
| SmsR6 |  | psRNA-78 | SMU_530c | 497858 | 498119 | SMU_531 | F | 262 |
| SmsR7 |  |  | SMU_1046c | 995311 | 995586 | SMU_1048 | R | 276 |
| SmsR8 | tmRNA |  | SMU_1196c | 1139336 | 1139677 | SMU_1197 | R | 342 |
| SmsR12 |  | psRNA-204<br>psRNA-205 | SMU_1862 | 1757885 | 1758495 | SMU_1865 | R | 611 |
| SmsR14 | 6S | psRNA-235 | SMU_2056 | 1929960 | 1930160 | SMU_2057c | F | 200 |
| SmsR16 |  |  | SMU_803c | 749489 | 749732 | SMU_804 | R | 244 |
| SmsR18 |  | psRNA-132 | SMU_1332c | 1254431 | 1255035 | SMU_1334 | R | 605 |
| SmsR19 |  |  | SMU_1398 | 1327742 | 1328337 | SMU_1400c | R | 596 |
| SmsR20 |  | psRNA-150 | SMU_1512 | 1439375 | 1439585 | SMU_1513 | R | 211 |
| SmsR22 |  |  | SMU_1703 | 1614938 | 1615314 | SMU_1704 | R | 377 |

<sup>a</sup>Small RNAs were named SmsR1-SmsR22 (Note: some of the original candidate sRNAs were later excluded from analysis).
<sup>b</sup>Existing Rfam entries for the putative sRNAs.
<sup>c</sup>Previously predicted sRNAs that overlap with our results (17).
<sup>d</sup>The putative 5' ends of sRNAs were estimated using visual scans of RNA-seq coverage plots for sites with sharp increases in reads mapped per nucleotide. 3' ends were estimated based on the locations of predicted terminators (**Table 2**) or for sRNAs without predicted terminators visual scans of the RNA-seq coverage plots were used to identify sites with sharp drops in reads per nucleotide. SmsR4 boundaries were confirmed using 5' and 3' RACE assays.

**Table 2.**
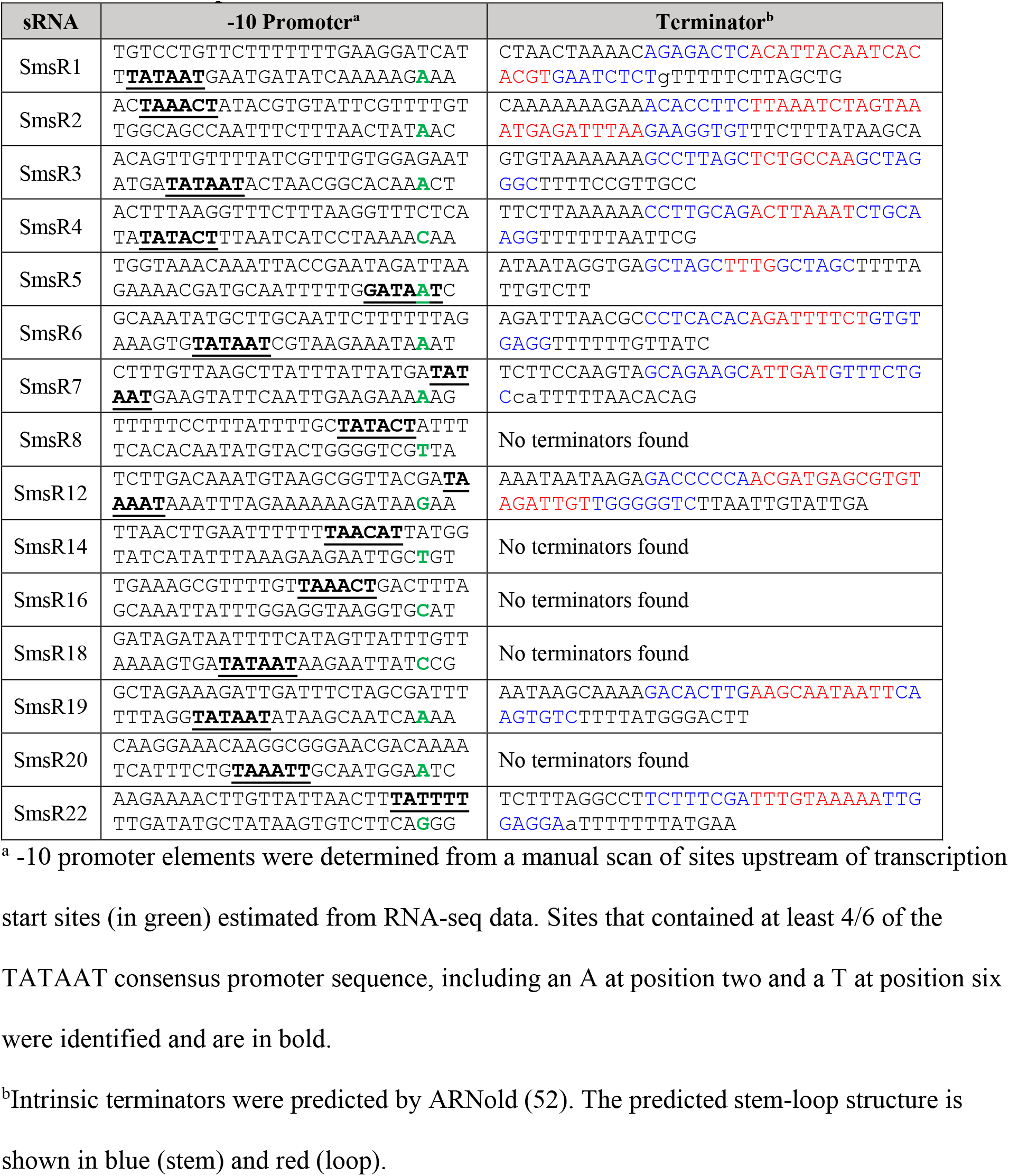
Predicted promoter and terminator elements of novel sRNAs.

### sRNAs respond to environmental stress

We evaluated the expression of the 15 novel sRNAs following *S. mutans* exposure to four stress-inducing conditions relevant to the human oral cavity: sugar-phosphate stress, hydrogen peroxide (H_2_O_2_), high temperature, and low pH (**Table 3**). All sRNAs showed patterns of differential expression in at least one of these conditions, suggesting likely roles in stress tolerance networks (**Figure 2**). Acid and sugar-phosphate stress induced differential expression of 12 sRNAs, while eight sRNAs were affected by heat stress and four responded to oxidative stress (**Figure 2**). Upregulation of sRNA expression was more common than downregulation across all conditions except for heat stress.

**Figure 2.**
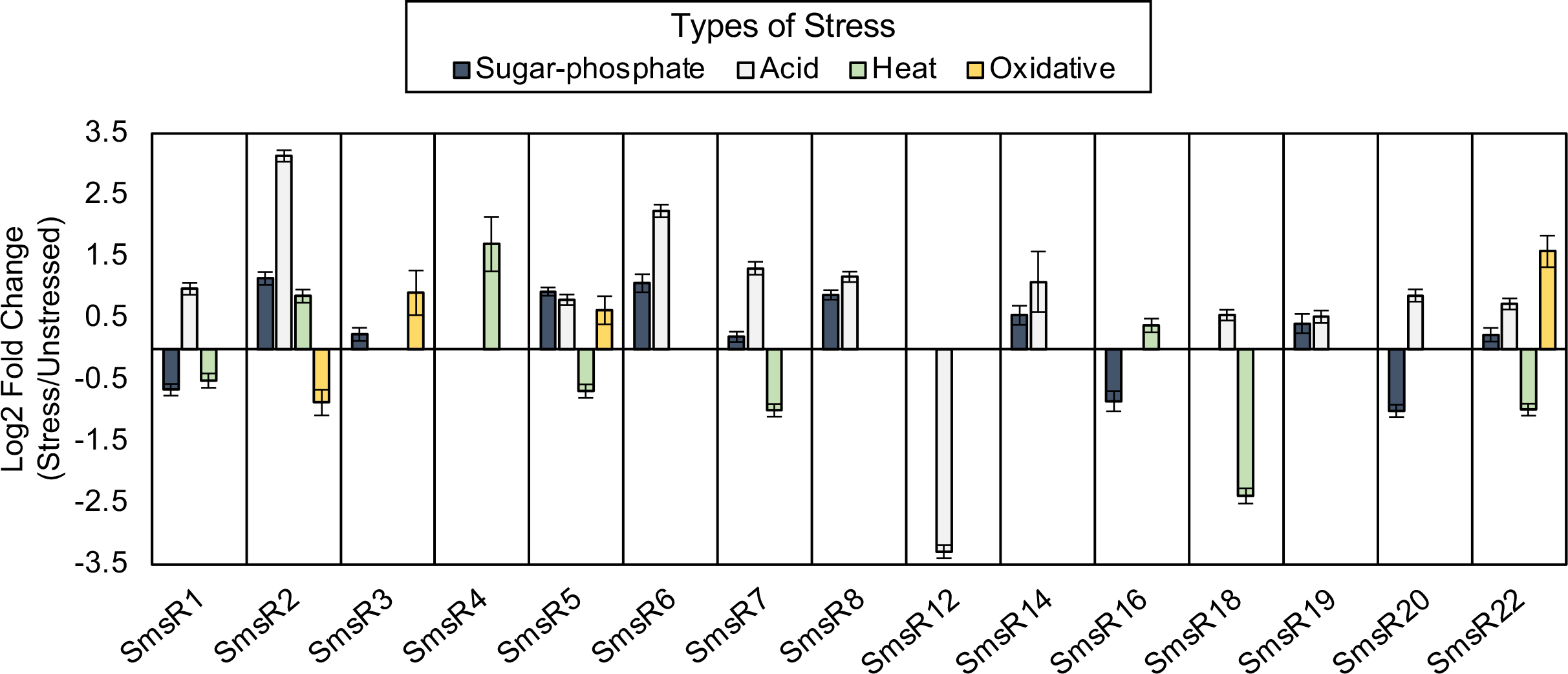
Differential expression of novel sRNAs in response to stress. Expression values for sRNAs were calculated from the total number of mapped RNA-seq reads. For each sRNA, Log2 fold change between treatment (stress) and control (no stress) samples (+/-standard error) from three replicate experiments are shown, except for acid stress, which used two replicates. Only expression values that were significantly different (p ≤ 0.05, Wald test) between stress and control conditions are displayed.

**Table 3.** Stress conditions tested in this study.

| Stress Type | Condition | Exposure Time |
| --- | --- | --- |
| Sugar-phosphate | 6% xylitol | 30 minutes |
| Oxidative | 1mM H<sub>2</sub>O<sub>2</sub> | 15 minutes |
| Heat | 45°C | 15 minutes |
| Acid | pH 5 | 30 minutes |

### SmsR14 and SmsR4 are unrelated sRNAs

6S RNA, a global regulator of transcription, is widely conserved among bacteria (24–26). Two copies of 6S RNA are present in *Bacillus subtilis*, a model gram-positive bacterium: one in the intergenic region between genes for azoreductase and RecQ helicase, and the other between two tRNA-associated genes (**Figure 3**) (27). The prevalence of 6S RNA in *Streptococcus* has not been examined previously. SmsR14, one of the newly-discovered sRNAs, is a 6S homolog and is located between *rarA* and tRNA-Lys genes in *S. mutans* (**Figure 3**). A covariance modeling (cm)-based search for 6S homologs in other *Streptococcus* species revealed only one copy of the sRNA in all genomes we analyzed (**Figure 4**). Interestingly, another novel sRNA (SmsR4) occupies the genomic location at which 6S RNA is present in *Escherichia coli*, i.e., upstream of the gene encoding 5-formyltetrahydrofolate cyclo-ligase (5-FTC) (**Figure 3**) (24, 26, 28).

**Figure 3.**
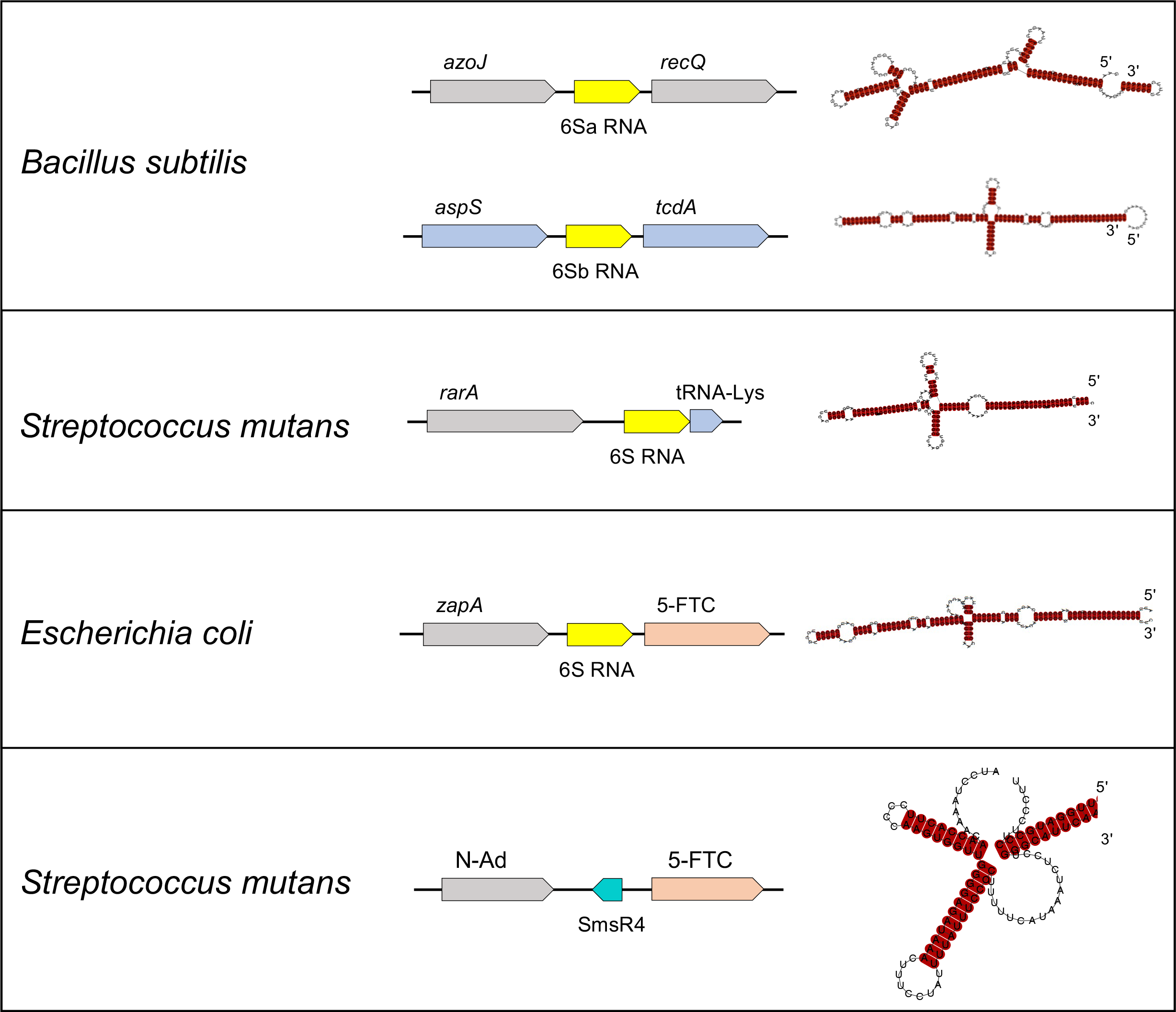
6S RNA and SmsR4 are distinct sRNAs. The genome locations and predicted secondary structures of 6S RNAs in *Bacillus subtilis*, *Streptococcus mutans,* and *Escherichia coli* are shown in the top three panels. The genome location and secondary structure of SmsR4 in *S. mutans* is displayed in the bottom panel. Genes that encode N-acetyldiaminopimelate deacetylase and 5-formyltetrahydrofolate cyclo-ligase are abbreviated as N-Ad and 5-FTC, respectively. (Note: genes are not drawn to scale.)

**Figure 4.**
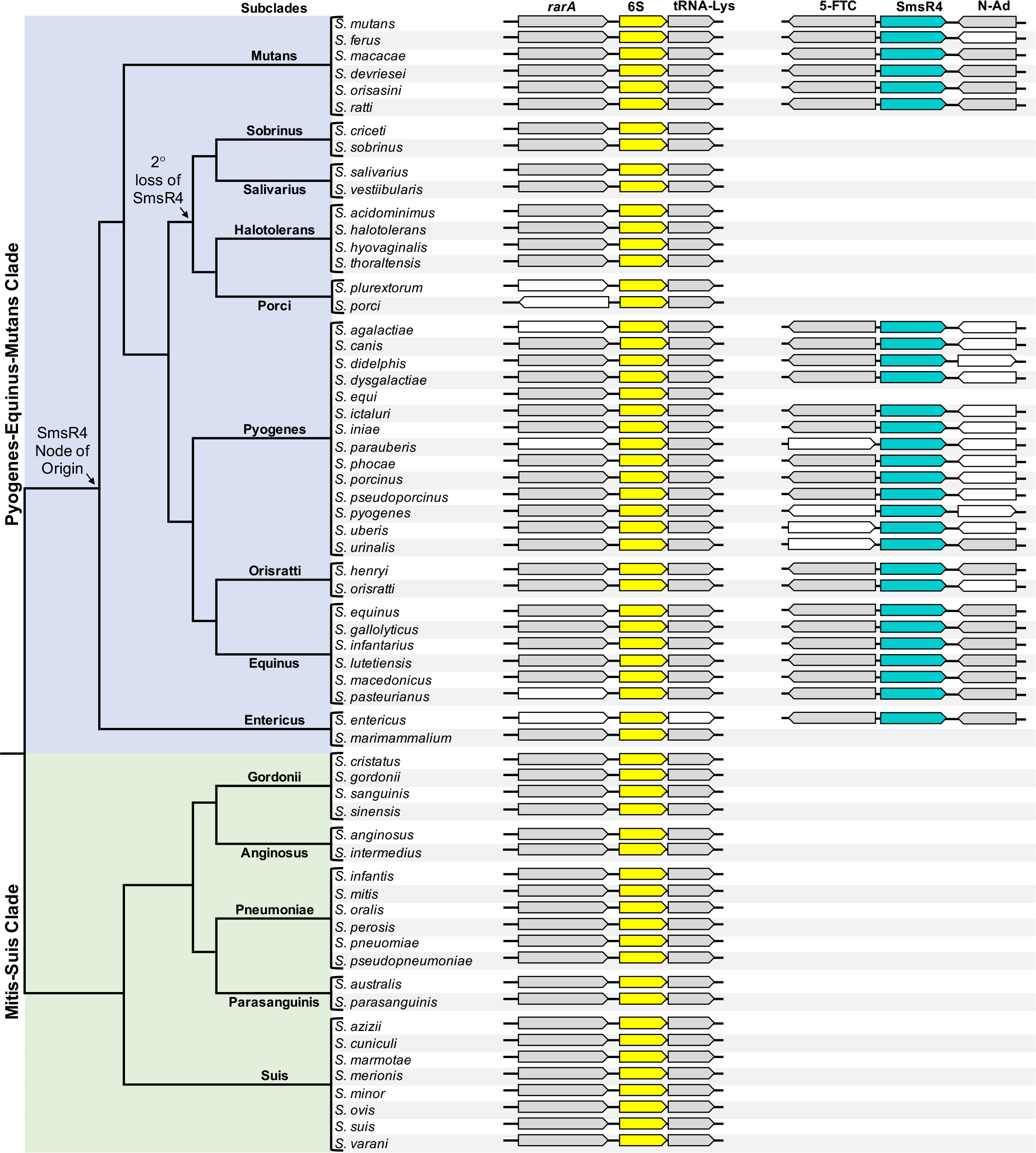
Prevalence of 6S RNA and SmsR4 in *Streptococcus*. Presence or absence of 6S and SmsR4 as determined by a covariance modeling-based search is shown. In most species, 6S is present between genes for RarA and tRNA-Lys, whereas SmsR4 is located between genes that encode 5-formyltetrahydrofolate cyclo-ligase (5-FTC) and N-acetyldiaminopimelate deacetylase (N-Ad). Flanking genes other than *rarA*, tRNA-Lys, 5-FTC, and N-Ad are shown as white arrows. The cladogram is from Patel and Gupta 2018 (29). (Note: genes are not drawn to scale.)

SmsR4, however, is much shorter than 6S (110 nt vs 194 nt), has a predicted secondary structure that is distinct from that of 6S, and is encoded on the opposite strand as 5-FTC gene (**Figure 3**). These differences suggest that SmsR4 is unrelated to 6S RNA and could have distinct functions in *S. mutans*.

### SmsR4 arose in the Pyogenes-Equinus-Mutans clade of *Streptococcus*

The genus *Streptococcus* could broadly be classified into two clades: Mitis-Suis and Pyogenes-Equinus-Mutans (29). Our cm-based search identified SmsR4 homologs only in the Pyogenes-Equinus-Mutans clade (**Figure 4**). Further, an evolutionary reconstruction showed that SmsR4 arose at the root of this clade but was later lost in the common ancestor of Sobrinus, Salivarus, Halotolerans and Porci subclades. While the rest of the Pyogenes subclade members contain SmsR4, it is absent in *Streptococcus equi*; similarly, in Entericus subclade*, Streptococcus marimmalium* has lost the sRNA, but it is retained by *Streptococcus entericus*. Despite these disparate cases of sRNA loss, the broad pattern of conservation of SmsR4 across a major clade of *Streptococcus* suggests that the novel sRNA has important functions.

### SmsR4 promotes *S. mutans* growth in sorbitol-containing media

To identify the functions of SmsR4 in *S. mutans*, we first confirmed the 5′ and 3′ boundaries of SmsR4 and generated an SmsR4-deletion (DEL) strain. We measured the growth of wild-type (WT) and DEL strains using a phenotypic microarray (30). In this analysis, DEL had reduced growth in comparison to WT in media containing sorbitol as the sole carbon source (**Figure S1**). We verified this phenotype by measuring bacterial growth in BTR medium that contained either 0.5% glucose (BTR-G) or 0.5% sorbitol (BTR-S). In this assay DEL had weaker growth than WT in BTR-S despite displaying a growth pattern identical to that of WT in BTR-G (**Figure 5**). The growth defect of DEL in BTR-S was overcome by a complementation strain (COMP) in which SmsR4 was expressed on the shuttle vector pDL278 (31, 32), indicating that the loss of SmsR4 caused DEL’s reduced growth. Probably because multiple copies of the SmsR4-carrying plasmid were present in each cell, COMP grew considerably better than WT in BTR-S, again denoting a role for the sRNA in promoting robust bacterial growth when using sorbitol as the carbon source.

**Figure 5.**
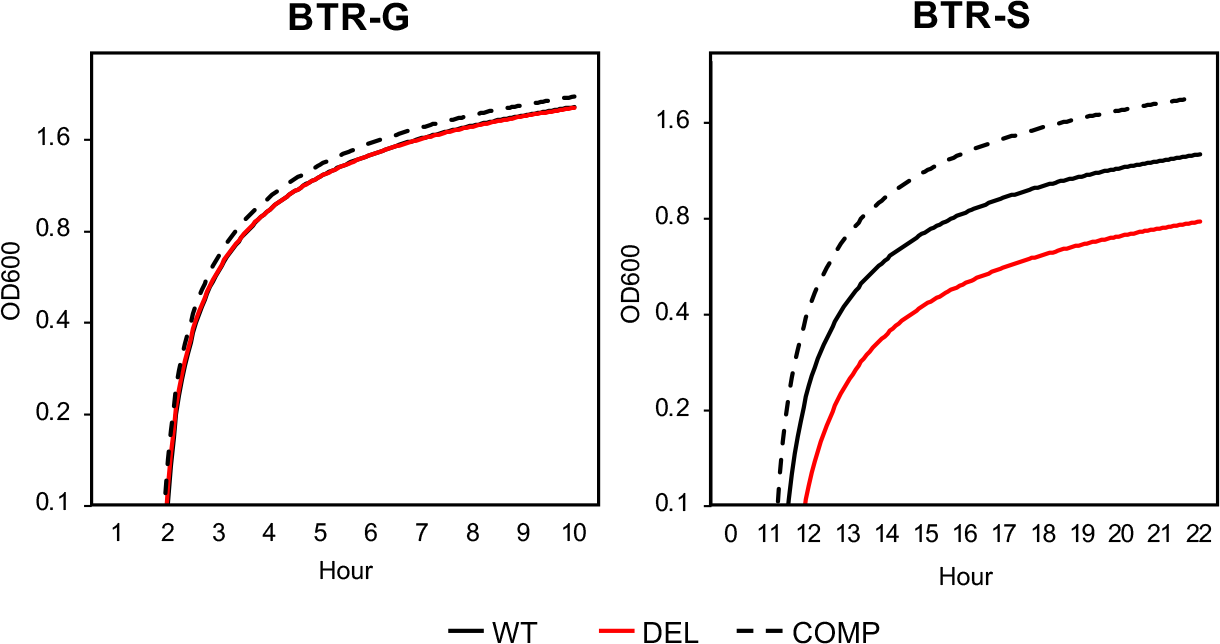
SmsR4 promotes *S. mutans* growth in sorbitol-containing medium. Growth of wild-type (WT), SmsR4-deletion (DEL), and SmsR4 complementation (COMP) strains in BTR medium with glucose (BTR-G, left) or sorbitol (BTR-S, right).

### SmsR4 regulates the EIIA component of the sorbitol PTS

In accordance with SmsR4’s potential role in regulating bacterial growth on sorbitol, *in silico* sRNA target prediction indicated that SmsR4 could bind to SMU_313, the gene encoding enzyme IIA of the sorbitol PTS (**Figure 6**; **Table S1)**. An SMU_313-deletion strain failed to achieve meaningful growth in BTR-S but grew at comparable levels to WT in BTR-G (**Figure S2**), demonstrating that SMU_313 is essential for growth on sorbitol but not glucose. We confirmed the interaction between SmsR4 and SMU_313 mRNA with an RNA-RNA electrophoretic mobility shift assay (EMSA), which showed that SmsR4 binds well to the 5′ UTR of SMU_313 mRNA (**Figure 7A**). The putative SmsR4-binding site identified by *in silico* analyses (**Figure 6**) is likely required for this interaction, as a mutation of the predicted SmsR4-binding site in SMU_313 inhibited its *in vitro* interaction with SmsR4 (**Figure 7B**). Taken together, our data indicate that SmsR4 modulates sorbitol import into the cell, likely by antagonizing translation of SMU_313 mRNA, which encodes the EIIA subunit of the sorbitol PTS.

**Figure 6.**
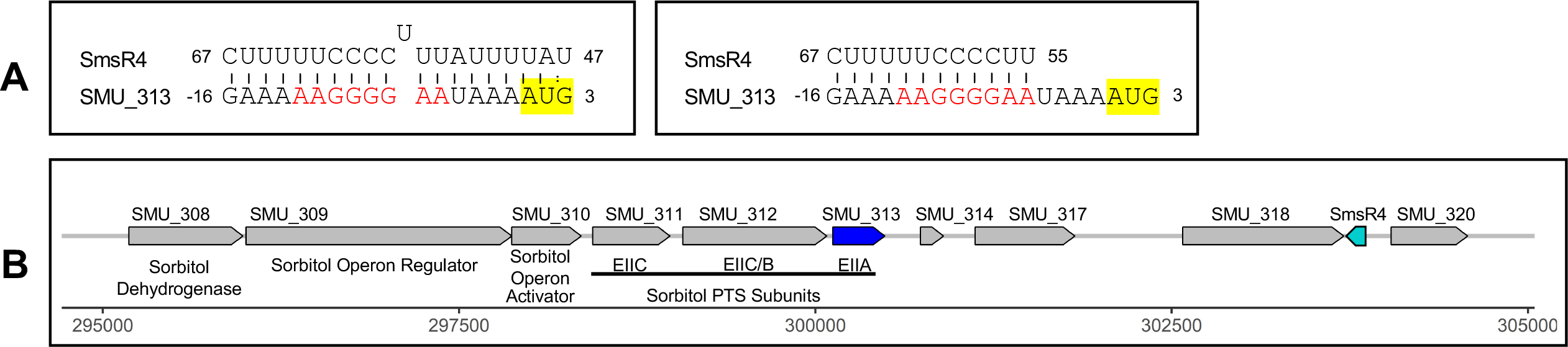
SmsR4 is predicted to bind to SMU_313. A) Two target prediction algorithms, IntaRNA (67) (left) and TargetRNA2 (68) (right), identified SMU_313 as a potential target of SmsR4. The predicted interaction sites on SmsR4 and 5′ untranslated region of SMU_313 are shown, and the start codon (AUG) of SMU_313 has been highlighted. The nucleotides highlighted in red are required for efficient binding of SmsR4 to SMU_313 (see **Figure 7**). **B)** Genomic locations of sorbitol phosphotransferase (PTS) system operon, including SMU_313 (blue), and SmsR4 (teal) in *S. mutans*.

**Figure 7.**
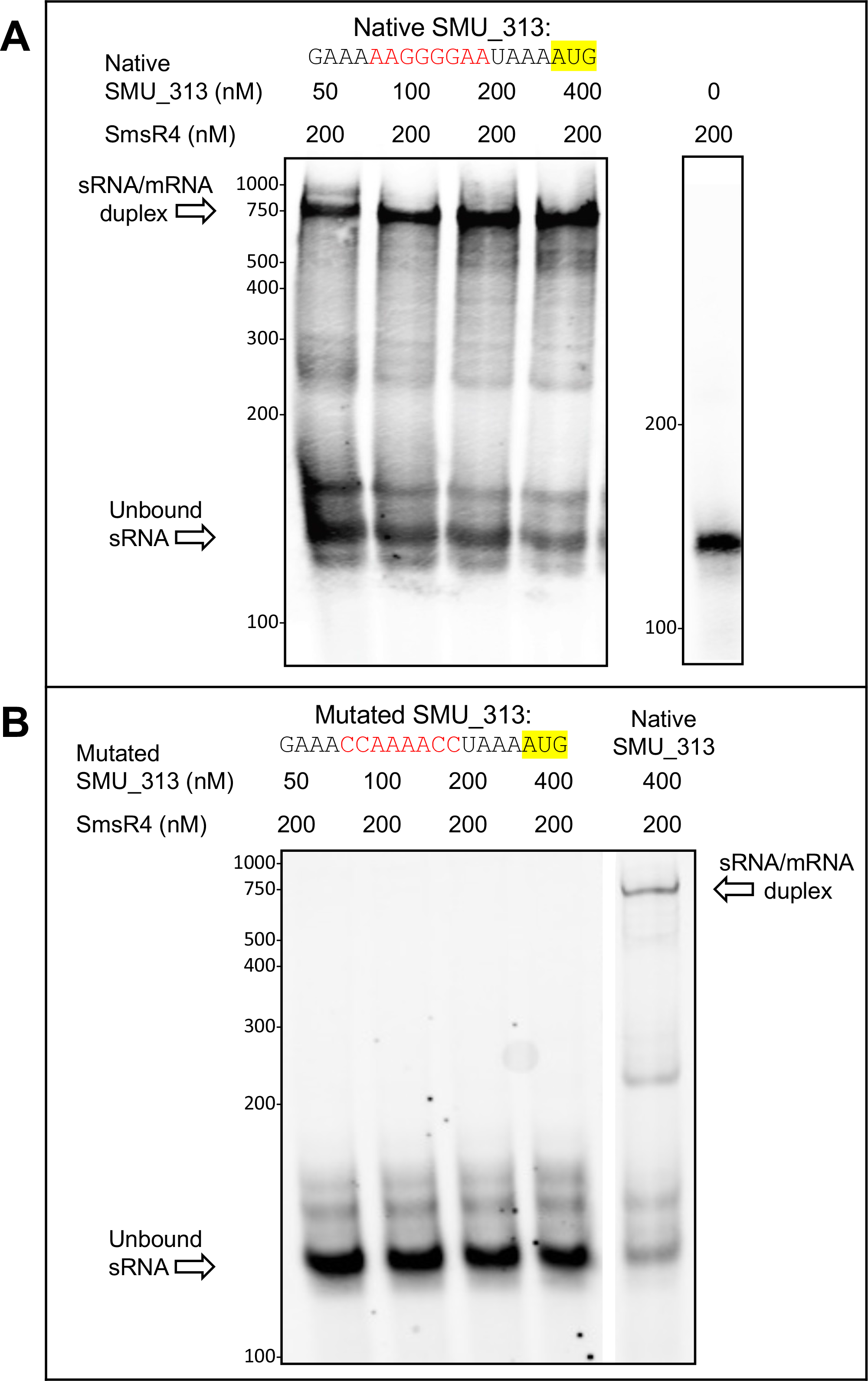
**SmsR4 binds to SMU_313**. **A)** An RNA-RNA electrophoretic mobility shift assay (EMSA) shows that increasing concentrations of SMU_313 with native 5′ untranslated sequence binds well to SmsR4. A control lane with unbound SmsR4 is shown to the right. **B)** SMU_313 transcript with a mutated binding site does not interact with SmsR4. A control lane with native SMU_313 transcript ran on the same gel is shown on the right. The native and mutated SmsR4-binding sites on SMU_313 are shown in red, and start codon is highlighted in yellow.

## DISCUSSION

sRNAs are critical for posttranscriptional gene regulation in bacteria, but their roles in the dental pathogen *S. mutans* have remained largely unknown. Prior to our study, a mostly bioinformatics-based analysis of gene expression in *S. mutans* grown with various carbon sources identified 243 sRNA candidates (17). In contrast, our approach combined genome-wide analysis with experimental validation and uncovered 15 sRNAs expressed under a single growth condition (BHI, OD_600_ 0.5-0.6) (**Figure 1**); hence, it is certainly possible that *S. mutans* transcribes additional undetected sRNAs in growth conditions different from those examined here. We noticed several sRNAs that appeared to be processed from larger parent transcripts, e.g., SmsR3 and SmsR20 (**Figure 1**). The generation of sRNAs from processed mRNAs has been well documented; for example, OppZ and CarZ are produced by RNase E cleavage of the *oppABCDF* and *carAB* mRNAs in *Vibrio cholera*, ArgX is produced from *argR* mRNA in *Lactococcus lactis*, and RsaC is formed via RNase III digestion of *mntACB* mRNA in *Staphylococcus aureus* (20, 33–37). Alternatively, some of these sRNAs may be regulatory elements with multiple products formed from premature transcriptional termination and transcriptional read-through (38).

The genomic contexts of sRNAs likely have implications for their functions (**Figure 1**). For instance, some of the sRNAs located in the 5′ UTRs of downstream genes (e.g., SmsR5, SmsR6, SmsSR8, SmsR16, SmsR20, SmsR22) could function as riboswitch-like elements (39), or sRNAs that are transcribed divergently from downstream protein-coding genes (e.g., SmsR1, SmsR3, SmsR19) could function as antisense sRNAs that regulate cis-encoded targets (40, 41). A few sRNA genes in other bacteria have been shown to contain small open reading frames (ORFs) that encode proteins (42). We searched the 15 sRNA sequences for ORFs and found that SmsR12 potentially encodes a small protein (33 amino acids). Although we could not detect a ribosome binding site upstream of the ORF and a BlastP search did not produce any hits in other bacteria, further studies would be required to determine whether the ORF is indeed functional.

Because sRNAs identified in this study exhibited differential expression under stress and their existence as discrete transcripts were validated via Northern blot, it is likely that they represent bona fide sRNAs participating in regulatory networks. Aside from SmsR4, the functions of these novel sRNAs are currently unknown, but preliminary *in silico* target predictions suggest that many of them are involved in processes critical to adaptation and virulence. For instance, SmsR2 and SmR22 are predicted to bind to PTS components, indicating that additional sRNAs may regulate sugar transport (**Table S1**), a process critical for *S. mutans* cariogenicity.

Among the 15 novel sRNAs, SmsR4 was intriguing because its genomic location is similar to that of 6S RNA in many proteobacteria (24), and as observed for 6S, SmsR4 was maximally expressed during the transition from exponential to stationary phase (**Figure S3**) (25). In gammaproteobacteria, expression of 6S is thought to be controlled by its linkage to the neighboring 5-FTC gene, which responds to nutrient limitation during the transition to stationary phase (24). In a similar manner, SMU_320, the 5-FTC homolog in *S. mutans,* could influence the expression of SmsR4; however, unlike with 5-FTC and 6S in gram-negative bacteria, SMU_320 and SmsR4 are encoded on opposite DNA strands and hence are not co-transcribed. Instead, a unique promoter likely initiates SmsR4 transcription (**Table 2**).

Because sugar transport is central to the proliferation and cariogenicity of *S. mutans*, it is unsurprising to find an sRNA regulator of its sorbitol usage. While further work is required to delineate the molecular details of its action, our results indicate that SmsR4 likely functions by modulating translation of the EIIA component of the sorbitol PTS. When sugars are transported through PTS systems, EIIA (e.g., SMU_313) participates in phosphorylation of the incoming sugar molecule as it enters into the cytoplasm (43). If left unchecked, phosphorylated sugars can accumulate in the cell and trigger sugar-phosphate stress (44). In *E. coli* and *Salmonella enterica,* the sRNA SgrS plays a key role in restoring glycolytic balance during sugar-phosphate stress by blocking the translation of mRNAs encoding the corresponding sugar-transporters (45, 46). In a similar fashion, SmsR4 could be modulating sorbitol intake by regulating the expression of SMU_313 (EIIA^sorb^) to relieve *S. mutans* sugar-phosphate stress during the transition from exponential to stationary phase growth in sorbitol-containing growth medium. Alternatively, modulation of SMU_313 expression by SmsR4 could impact alternative functions of EIIA. For example, this protein has been shown to negatively impact glycerol metabolism in *Klebsiella pneumoniae* and to reduce *S. enterica* virulence (47, 48). In conclusion, our *in silico* and biochemical data both support a role for sRNAs as posttranscriptional regulators of sugar transport through PTS systems. Further insights into these regulatory mechanisms may provide new opportunities to develop specific inhibitors of *S. mutans* growth in the oral cavity.

## MATERIALS AND METHODS

### Bacterial strains and growth assays

Growth experiments were conducted by diluting overnight cultures of *S. mutans* UA159 in fresh media. BTR broth base (1% Bacto-Tryptone, 0.1% yeast extract, 0.61% K_2_HPO_4_, 0.2% KH_2_PO_4_) supplemented with either 0.5% glucose (BTR-G) or 0.5% sorbitol (BTR-S) was utilized for growth assays. For experiments comparing the growth of WT, DEL, and COMP strains, 1 mg/ml spectinomycin was added to the growth media to retain the pDL278 plasmid. All growth assays were done at 37°C in an anaerobic chamber (5% hydrogen, 5% carbon dioxide, 90% nitrogen).

### Phenotypic microarray

Biolog Phenotypic microarray assays were conducted per manufacturer’s recommendations (36). Briefly, overnight cultures from single bacterial colonies were diluted in fresh Brain Heart Infusion (BHI) broth and grown to an OD_600_ of 0.6-0.7 at 37°C in an anaerobic chamber. Cells were collected via centrifugation (3,000xg, 5 min) and washed twice with PBS and resuspended to an OD_600_ of 0.4 in IF-0a GN/GP base. Inoculating fluid was prepared and combined with cells at 81% turbidity and 100 μl of the mixture was added to each well and overlaid with 40 μl mineral oil. Temperature was maintained at 37°C while absorbance values at 590 nm and 750 nm were collected every 20 min for 24 h using a Multiskan Spectrum plate reader (Thermo Fischer Scientific). Results were obtained by subtracting the measurements at 750 nm from those at 590 nm, and the average of two replicates was used to construct a growth curve for each well.

### sRNA discovery

Bacterial cultures were grown in BHI broth to an OD_600_ of 0.5-0.6. RNA stop solution (5% Phenol, 95% Ethanol) was added to bacterial cultures (1.25 mL stop solution per 10 mL culture) and cells were pelleted by centrifugation at 10,000xg for 10 min at 4°C. Bacterial pellets were resuspended in 1 ml of TRI reagent (Thermo Fischer Scientific) and total RNA was extracted using the manufacturer’s protocol. RNA was resuspended in nuclease-free water and DNA was removed by TURBO DNase (Thermo Fischer Scientific) treatment. RNA sequencing was performed at the Yale Center for Genome Analysis using Illumina NovaSeq (paired-end, 150 bp). Raw reads have been deposited in the NCBI Sequence Read Archive under the accession number PRJNA726731. RNA-seq reads were processed using Trimmomatic to remove low-quality reads and adapters (49). CLC Genomics workbench was used to map the reads to the *S. mutans* UA159 genome (NC_004350.2) and to determine the total read count for each gene.

Coverage plots were generated by calculating reads mapped per nucleotide across the entire genome using an in-house Perl script, as described previously (13). The Artemis genome browser (50) was used for visual inspection of transcriptomics data and gggenes package in R (version 0.4.1) was used to draw Figure 1. The putative 5′ end of each sRNA was estimated from sites on transcriptional coverage plots with sharp increases in reads mapped per nucleotide. RNAalifold was used to predict secondary structure of SmsR4 and 6S RNA (51), and ARNold was used to predict 3′ Rho-independent terminator structures (52). For sRNAs without predicted terminators, 3′ ends were defined as sites on transcriptional coverage plots with a sharp decrease in reads mapped per nucleotide. Potential open reading frames (ORFs) within sRNA genes were detected using ORF Finder (53).

### Stress induction

Overnight cultures of *S. mutans* UA159 were diluted 1:100 in BTR-G and grown to an OD_600_ of 0.3-0.4 at 37°C in an anaerobic chamber. To induce sugar-phosphate stress, 6% xylitol in BTR broth was added to one half (treatment) and grown for 15 minutes at 37°C, while an equal volume of BTR broth without xylitol was added to the control and incubated under the same conditions. Similarly, for oxidative stress induction, 1mM H_2_O_2_ was added to one half of the culture (treatment) while an equal volume of water was added to the control half and incubated for 15 minutes. For heat stress, one half of the culture was incubated at 37°C for 15 minutes (control), while the other was incubated at 45°C for 15 minutes (treatment). To induce acid stress, cultures were centrifuged and one half was resuspended in BTR-G, pH 7 (control), while the other half was resuspended in BTR-G, pH 5 (treatment), and incubated for 30 minutes at 37°C. Anaerobic conditions were maintained throughout the stress induction assays. RNA was extracted as described above. To confirm stress induction, qRT-PCR was used to measure the expression of stress marker genes previously associated with each tested stress condition (54–57) (**Figure S4**). Contaminating DNA was removed by TURBO DNase (Thermo Fischer Scientific) treatment and cDNA was synthesized with a High-Capacity cDNA Reverse Transcription Kit (Thermo Fischer Scientific) using random primers. qRT-PCR was performed using SYBR Green master mix (Thermo Fischer Scientific) and gene-specific primers (**Table S2**). RNA was sequenced and RNA-seq reads were processed as described above. The DESeq2 package in R was used to determine differential gene expression of sRNAs under the four stress-inducing conditions compared to controls (58). Experiments were performed in triplicate for all conditions except for acid stress, which was performed in duplicate.

### *In vitro* transcription

Amplification of gDNA for *in vitro* transcription was performed using PCR primers designed to incorporate a T7 promoter (**Table S2**). PCR products to be used as DNA templates were cleaned using a NucleoSpin Gel and PCR Clean-up kit (Takara Bio). *In vitro* transcription was performed using MAXIscript T7 Transcription Kit (Thermo Fischer Scientific) per manufacturer’s protocol with a maximum of 1 µg of DNA used as template. Following TURBO DNase treatment, RNA was purified using a Monarch RNA Cleanup Kit (New England Biolabs).

### Northern blot

RNA was isolated from *S. mutans* cells under sugar-phosphate stress, oxidative, heat, or acid stress as described above. Equal amounts of RNA were adjusted to 10 μl with nuclease-free water, and 10 μl of 2x RNA loading dye (Thermo Fischer Scientific) was added and incubated for 10 min at 70°C followed by 3 min on ice. Samples were loaded onto either 6% or 10% TBE-Urea Gel (Thermo Fischer Scientific) along with a biotinylated sRNA ladder (Kerafast). Gels were run in 1x TBE buffer at 180V for 60 min (6% gels) or 180V for 80 min (10% gels). RNA was transferred to a Biodyne B Nylon Membrane (Thermo Fischer Scientific) using the BioRad Mini-Trans Blot at 12V overnight, 4°C in 0.5x TBE buffer. Membranes were UV-crosslinked using a Staralinker 2400 UV Crosslinker (1200 mJ) and were moved to glass hybridization chambers and prehybridized using 10 ml of ULTRAhyb-Oligo Buffer (Thermo Fischer Scientific) at 45°C for 2 h with rotation. RNA probes produced from *in vitro* transcription as described above were heated at 95°C for 5 min and cooled on ice for 3 min, then added to fresh hybridization buffer. Membranes were incubated overnight at 45°C with rotation. After washing, membranes were incubated for 2 h with shaking in Licor Intercept Blocking Buffer with 1% SDS at room temperature. Blocking buffer was removed and membranes were incubated in Streptavidin-IRDye 800 CW diluted 1:20,000 in Licor Intercept Blocking Buffer with 1% SDS for 30 min. Blots were washed and viewed on a Licor Odyssey scanner.

### EMSA and mutagenesis

Electromobility shift assay (EMSA) was performed as described previously (59). Briefly, a DNA template was amplified from *S. mutans* UA159 gDNA using primers with a T7 tag (**Table S2**). RNA was transcribed from this template using the MAXI T7 Transcription Kit that incorporated biotinylated uracil into the sRNA transcript. RNA was purified with a Monarch RNA Cleanup Kit and resuspended in TE buffer. SmsR4 and SMU_313 transcripts were combined at ratios shown in **Figure 7** and heated for 5 min at 85°C, then immersed in ice for 30 sec. The reaction volume was adjusted to 10 μl with 5x TMN buffer and incubated at 37°C for 30 min. Samples were run on an 8% TBE gel (Thermo Fischer Scientific) for 90 min at 100 V in 1x TBE buffer. The gel was transferred to a Biodyne B Membrane overnight at 12V, 4°C in 0.5x TBE. Membranes were crosslinked, blocked, and probed with Strepatvidin-IRDye 800CW as described above for Northern blot assays, and images were examined on a Licor Odyssey scanner. SMU_313 with mutated SmsR4-binding site was constructed using the Q5 Mutagenesis Kit (New England Biolabs) and mutations were confirmed through Sanger sequencing.

### Gene deletion and complementation

SmsR4- and SMU_313-deletion strains were constructed using the markerless-mutagenic system and an IDFC2 selection and counter-selection cassette, as described previously (60). Complementation was preformed using the pDL278 plasmid designed for expression in both *E. coli* and *S. mutans* (31, 32). The plasmid was purified from *E. coli* using a Plasmid MiniPrep kit (Thermo Fischer Scientific). pDL278 was linearized using BamHI and EcoRI, and PCR products (**Table S2**) were ligated into the linearized plasmid using T4 ligase (Thermo Fischer Scientific). Plasmids were then transformed into competent *E. coli* DH5-alpha cells following manufacturer’s protocol (New England Biolabs). Sanger sequencing was used to confirm plasmid construction. Purified plasmids were transformed into *S. mutans* using competence stimulating peptide as described previously (60).

### RACE assay

Rapid Amplification of cDNA Ends (RACE) was performed using a RACE kit (Thermo Fisher Scientific) per the manufacturer’s recommendations. 5′ RACE assay was conducted using a gene specific primer (GSP) complementary to the 3′ end of SmsR4 (**Table S2**). RNA degradation, cDNA synthesis, and TdT tailing were accomplished using kit components and manufacturer protocols. A second nested GSP was used to amplify the tailed cDNA, and the PCR product from this reaction was cloned into pGEM T-Easy vector (Promega), transformed into competent DH5-alpha *E. coli* and the 5′ end of SmsR4 was determined using Sanger sequencing. For the 3′ RACE assay, Poly-A polymerase (New England Biolabs) was used to add poly-A tails to all transcripts. Precipitated poly-A tailed RNA was reverse transcribed using an oligo-dT adapter primer and SuperScript II reverse transcriptase. RNA was subsequently degraded using RNase H. A GSP designed to bind to the 5′ end of SmsR4 and an Abridged Universal Amplification Primer (**Table S2**) were used to amplify the cDNA, and PCR products were cloned into pGEM T-Easy vector, transformed into competent *E. coli* DH5-alpha cells and the 3′ end of SmsR4 was identified using Sanger sequencing.

### Covariance modeling

Covariance models of sRNAs were constructed as previously described (61, 62). Briefly, SmsR4 and 6S sequences from *S. mutans* UA159 were used as queries in BlastN searches against all *Streptococcus* genomes available in RefSeq (63, 64). Hits with >65% identity and >70% coverage were retained and five and six sequences were randomly selected to serve as seed sequences for constructing 6S and SmsR4 covariance models, respectively (**Table S3**). The WAR webserver was used to align the seed sequences and the Infernal suite of tools (v1.1.2) was used to construct (cmbuild) and calibrate (cmcalibrate) an initial covariance model for each sRNA (65, 66). This model was used to search (cmsearch) a database constructed from 37 representative *Streptococcus* full genomes available on RefSeq (**Table S4**). Results from cmsearch with an e-value <1e-5 were used to add unrepresented sequences to the query model, which was then refined, recalibrated, and used for another round of cmsearch. This process was repeated for each sRNA until cmsearch failed to yield new unrepresented sequences. The final models were used to determine the prevalence of 6S and SmsR4 in 62 full and partial *Streptococcus* genomes (**Table S5**). The presence or absence of each sRNAs was mapped on a previously published phylogenetic tree (29) and nodes of sRNA origin and secondary loss were determined through maximum parsimony.

### *In silico* sRNA target prediction

For SmsR4 target prediction using IntaRNA (67), SmsR4 sequence as determined by RACE assay was used as input and searched against the *S. mutans* UA159 genome using default parameters (75 nt upstream and downstream from the translation start site, one interaction per pair, 7 nt hybridization seed). Target RNA2 (68) was run using the same query sequence and default parameters (80 nt upstream and 20 nt downstream from the translation start site, 7 nt hybridization seed). IntaRNA was also used to predict targets for 11 other sRNAs using stricter parameters to identify potential interactions adjacent to translation start sites of mRNA targets. For all *in silico* target predictions, only significant results (p < 0.05, as determined by IntaRNA or Target RNA2) were retained.

## ACKNOWLEDGEMENTS

We thank Zhengzhong Zou and Samantha Fancher for their assistance with experiments, and Shaun Wachter for technical advice. This work was supported in part by National Institute of Dental and Craniofacial Research grants DE028409 to RR and DE028252 to JM.

## SUPPLEMENTAL FIGURE LEGENDS

**Figure S1. Phenotypic microarray.** Growth of wild-type (WT) and SmsR4-deletion (DEL) strains in Biolog medium with sorbitol as sole carbon source (30). Values represent the average of two independent growth experiments.

**Figure S2. Effect of SMU_313 deletion on growth in sorbitol.** Growth of SMU_313-deletion (SMU_313 DEL) and wild-type (WT) strains of *S. mutans* in BTR medium with glucose (BTR-G, left), or sorbitol (BTR-S, right).

**Figure S3. SmsR4 expression over time.** Northern blot for SmsR4 in wild-type *S. mutans* grown in BTR-S was performed at time-points shown. 5S RNA was used as a loading control.

**Figure S4. Confirmation of stress induction.** Either upregulation or downregulation of genes known to be associated with each stress condition were used to confirm stress induction (54–57). Values represent means (+/-standard error) from three independent qPCR assays, except for *groES* and SMU_1805, which were from two replicates.

## SUPPLEMENTAL TABLES

**Table S1.** Targets predicted by IntaRNA for novel sRNAs.

**Table S2.** Primers used in this study.

**Table S3.** Genomes used for initial SmsR4 and 6S covariance model construction.

**Table S4.** Genomes used for covariance model calibration.

**Table S5.** Genomes used to determine the prevalence of 6S and SmsR4 in *Streptococcus*.

